# Harnessing the endogenous Type I-C CRISPR-Cas system for genome editing in *Bifidobacterium breve*

**DOI:** 10.1101/2023.09.25.559376

**Authors:** Xiao Han, Lulu Chang, Haiqin Chen, Jianxin Zhao, Fengwei Tian, R. Paul Ross, Catherine Stanton, Douwe van Sinderen, Wei Chen, Bo Yang

## Abstract

*Bifidobacterium breve*, one of the main bifidobacterial species colonizing the human gastrointestinal tract in early life, has received extensive attention for its purported beneficial effects on human health. However, exploration of the mode of action of such beneficial effects exerted by *B. breve* is cumbersome due to the lack of effective genetic tools, which limits its synthetic biology application. Given the widespread presence of endogenous CRISPR-Cas systems in *B. breve*, the current study developed an endogenous CRISPR-based gene editing toolkit for genetic manipulation of *B. breve*. Deletion of the gene coding uracil phosphoribosyl-transferase (*upp*) was achieved in two different *B. breve* strains using this system. In addition, translational termination of uracil phosphoribosyl-transferase was successfully achieved in *B. breve* FJSWX38M7 by single-base substitution of the *upp* gene and insertion of three stop codons. The gene encoding linoleic acid isomerase (*bbi*) in *B. breve*, being a characteristic trait, was deleted after plasmid curing, which rendered it unable to convert linoleic acid into conjugated linoleic acid, demonstrating the feasibility of successive editing. This study expanded the gene manipulation toolkit of *B. breve* and provides a reference for functional genome editing and analysis using an endogenous CRISPR-Cas system in *Bifidobacterium*.

**Importance:** The lack of effective genetic tools for *Bifidobacterium breve* is an obstacle to studying the molecular mechanisms of its health-promoting effects, hindering the development of next-generation probiotics. Here, we introduce a gene editing method based on the endogenous CRISPR-Cas system, which can achieve gene deletion, single base substitution, gene insertion and continuous gene editing in *B. breve*. This study will promote the excavation of functional genes and elucidation of molecular mechanisms of *B. breve*.

## Introduction

Bifidobacteria are commensal bacteria that colonize the human gut early in life, in some cases constituting more than 90% of the total bacterial population in a given individual (1, 2). Certain *Bifidobacterium* strains have been added to functional foods as bioactive ingredients due to their purported benefits on human health or have been commercially used as adjuvants for drug therapy (2). *Bifidobacterium breve*, a common and abundant component of the gut microbiota in breastfed newborns, has been associated with various beneficial or probiotic activities, such as its ability to alleviate gastrointestinal disorders and its contribution to the maturation of the immune system in infants (3). Although various health benefits have been attributed to *B. breve*, the molecular mechanisms supporting these probiotic activities are poorly understood, which has impeded their regulatory acceptance as probiotics in the European Union and other jurisdictions, while hindering future development and discovery of new functions. In fact, genetic manipulation of *B. breve* can be challenging due to its thick cell walls, oxygen sensitivity, active restriction modification (R-M) systems, and lack of genetic toolkits (4). The current method commonly used for gene inactivation in *B. breve* is based on homologous recombination using a non-replicating plasmid due to the lack of a functional replicase-encoding *repA* gene, or by transposon mutagenesis (5–10). But these approaches typically introduce antibiotic resistance genes into the genome, not allowing for successive gene editing. Moreover, low transformation frequency and/or low frequency of transposition or homologous recombination events may make it difficult to reliably generate mutants in many strains (11).

The clustered regularly interspaced short palindromic repeats (CRISPR)-CRISPR-associated proteins (Cas) system is one of the immune systems of bacteria against foreign DNA invasion and has been designed for genome editing in a variety of organisms (12). CRISPR-Cas9-based gene editing in bacteria was initially performed in *Escherichia coli* and subsequently realized in other species (13, 14). However, the large size of the gene encoding spCas9 (from *Streptococcus pyogenes*, 1368 AA) reduces the transformation efficiency of a plasmid carrying this gene, and prevents its efficient delivery to certain bacteria (15). Some studies have shown that SpCas9 may be cytotoxic, leading to abnormal cell morphology and reduced colonies (16, 17). Genome editing using a native CRISPR-Cas system is a potentially less harmful approach that has been successfully applied in *Zymomonas mobilis* (18), *Lactobacillus crispatus* (19) and *B. animalis* subsp. *lactis* (20). Since its effector protein cascade is expressed by the strain itself, the size of the plasmid can be greatly reduced to ensure transformation efficiency. In our previous study, the types of CRISPR-Cas systems and CRISPR loci in *B. breve* were characterized (21). The CRISPR-Cas system widely exists among *B. breve* strains (47%), among which Type I-C accounts for the highest proportion. The widely distributed CRISPR-Cas system provides the basis for the development of an autochthonous CRISPR-based gene editing toolkit.

In this study, we first characterized the activity of the Type I-C CRISPR-Cas system in *B. breve*. An endogenous CRISPR-based gene editing toolkit was then established by designing a plasmid containing an artificial crRNA and a repair template. Then gene deletions, single-base substitutions, short DNA insertions, and successive gene editing were performed in *B. breve* using this toolkit. This work thus provides a molecular tool for exploring the molecular mechanism of the functional activity of *B. breve*, while it furthermore represents a reference for the application of endogenous CRISPR-Cas systems in other bacteria.

## Results

### Investigating the activity of an endogenous CRISPR-Cas system in *B. breve*

According to our previous research, about 47% of *B. breve* strains encompass an apparently intact, endogenous CRISPR-Cas system, among which Type I-C was shown to be the most prevalent (21). The Cas5, Cas7, and Cas8 proteins constitute a Type I-C cascade, in which Cas5 is responsible for crRNA processing (22). Following recognition of the protospacer adjacent motif (PAM) sequence by the cascade, crRNA base pairs with the target DNA strand to induce R-loop formation. The cascade then recruits Cas3, which initiates degradation of the non-target strand (23) (Fig. S1). A Type I-C CRISPR-Cas system was identified in *B. breve* FJSWX38M7, consisting of a CRISPR array (82 spacers) and a *cas* gene cluster (Fig. 1A). In order to investigate the activity of the Type I-C system in FJSWX38M7, a plasmid interference assay was designed. The *E. coli*/*Bifidobacterium* shuttle vector pNZ123, carrying a chloramphenicol resistance gene, was used as a framework for the plasmid interference assay. The PAM sequence was predicted to be 5’-TTC-3’ in the *B. breve* Type I-C system based on our previous study (21). We identified 81,083 instances of the 5′-TTC-3′ or 5′-GAA-3′ motifs in the *B. breve* FJSWX38M7 genome. The number of nucleotides between two neighboring PAM sequences was also counted, with an average of about 64 nucleotides presenting a PAM motif, allowing for a wide range of genetic manipulations (Fig. S2). The last obtained spacer in the FJSWX38M7 CRISPR array was cloned into pNZ123 with or without a PAM sequence, generating plasmids pNZ123-PS82 and pNZ123-S82, respectively. Compared with pNZ123-S82, the transformation efficiency of pNZ123-PS82 was reduced by more than four orders of magnitude, strongly implying that the active CRISPR-Cas system created a barrier to the entry of exogenous DNA with PAM and spacer.

**Fig. 1.**
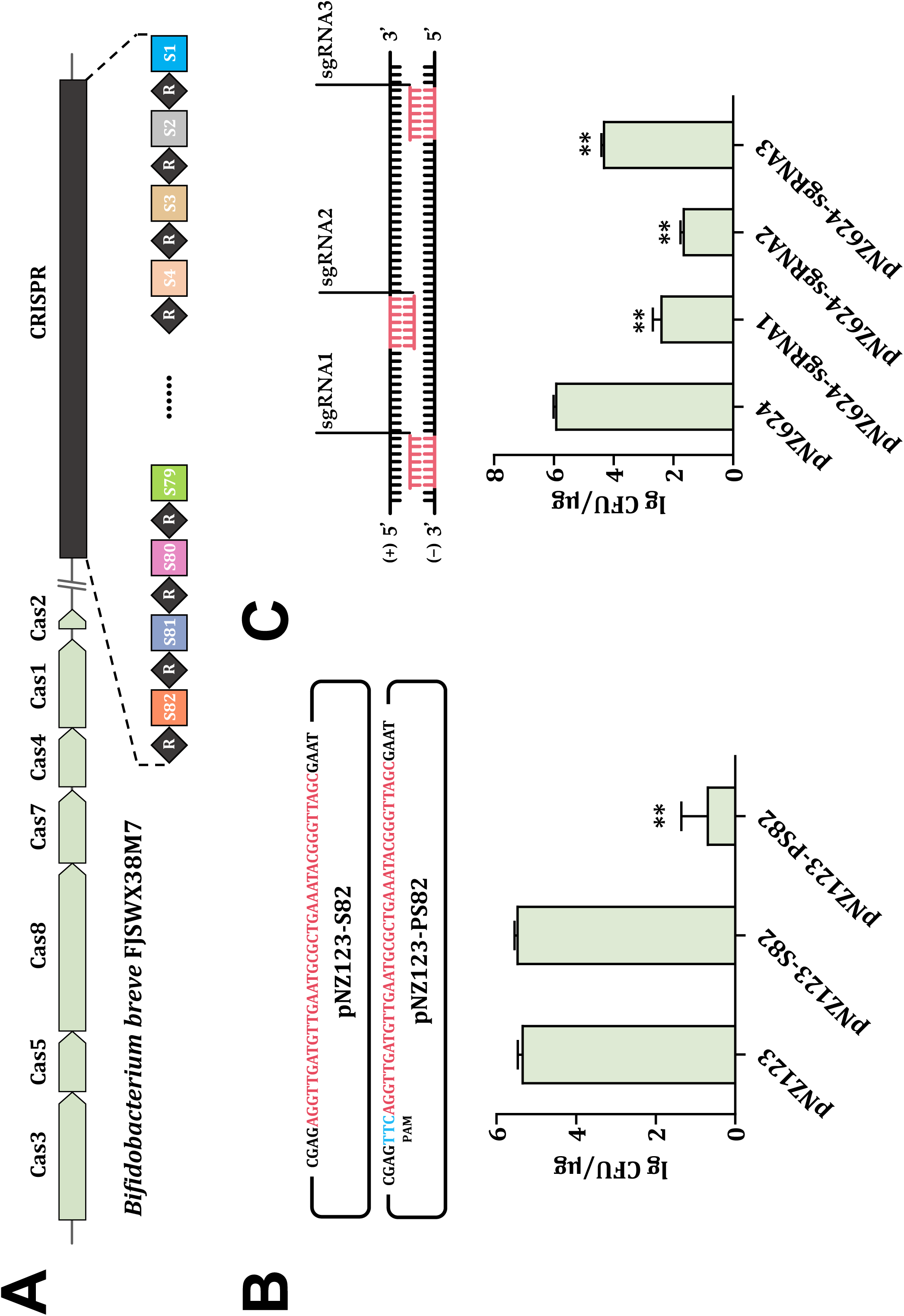
Endogenous Type I-C CRISPR locus in *B. breve* FJSWX38M7 and its activity. (A) Schematic representation of the CRISPR array and *cas* loci in *B. breve* FJSWX38M7. The *cas* genes are indicated by green arrows. Repeats are indicated by black diamonds. Each differently colored square represents a unique spacer, and the first spacer acquired named S1 is shown on the right. (B) Plasmid interference assay. pNZ123-PS82 contains the spacer 82 and PAM sequence in FJSWX38M7 CRISPR array, while pNZ123-S82 only contains spacer 82 without the PAM sequence. The transformation efficiencies of these plasmids were represented by log_10_ CFU/μg. The values were mean ± SD of three independent experiments. The empty plasmid pNZ123 was used as control. \*\**P* < 0.01 was determined by an unpaired *t*-test. (C) Self-target assay. pNZ624-sgRNA1-3 contained three different sgRNAs targeting the *upp* gene, respectively. sgRNA2 targeted the forward strand and sgRNA1/sgRNA3 targeted the reverse strand. The empty plasmid pNZ624 was used as a control. \*\**P* < 0.01 was determined by an unpaired *t*-test.

After the activity of endogenous I-C CRISPR-Cas was demonstrated in *B. breve* FJSWX38M7, an artificial CRISPR crRNA designed to target arbitrary genes was constructed for gene editing by repurposing the endogenous system. The artificial crRNA included a leader sequence, a sgRNA insertion site, two repeats, and a rho-terminator. The native leader of the I-C CRISPR array in *B. breve* was first analyzed, removing strains whose leader sequence was interrupted by mobile elements (Fig. S3). The results showed that the leader sequence was relatively conserved, and the nucleic acid with the highest relative frequency at each position was selected to form the final leader sequence. The repeat sequence was predicted using CRISPRCasFinder (https://crisprcas.i2bc.paris-saclay.fr/CrisprCasFinder/Index) (24), and a short sequence with two BsaI sites was designed in the middle of the two repeat sequences to allow for convenient sgRNA insertion (Fig. S4). Synthetic artificial crRNA was cloned into pNZ123 to generate the gene editing plasmid pNZ624. The self-targeting experiment was designed to verify the effect of artificial crRNA. The number of different spacer lengths in the *B. breve* I-C CRISPR-Cas system was analyzed. The vast majority of spacers were 35 nucleotides long, which was a reference for the length design of sgRNA (Fig. S5). Three different sgRNAs were designed based on the *upp* gene as the target gene, among which sgRNA1 and sgRNA3 targeting the non-coding strand, and sgRNA2 targeting the coding strand. Three sgRNAs were ligated to BsaI-digested pNZ624 to generate pNZ624-sgRNA1, pNZ624-sgRNA2, and pNZ624-sgRNA1, respectively. The transformation efficiency of FJSWX38M7 decreased significantly when using these three plasmids when compared with pNZ624, in particular the transformation efficiency of pNZ624-sgRNA2 was reduced by more than 10,000-fold (Fig. 1C). These results clearly indicate that the nuclease Cas3 of the strain cuts its own genome, being under the guidance of the cascade containing sgRNA, thus leading to cell death, which demonstrated the potential of the endogenous CRISPR-Cas system in gene editing.

### Establishment of the Type I-C CRISPR-based gene deletion toolkit for *B. breve*

DNA single-strand breaks (SSBs) or double-strand breaks (DSBs) pose a deadly threat to bacteria. Different from eukaryotes, which often use non-homologous end joining (NHEJ) to repair DNA damage, NHEJ is not ubiquitous in prokaryotes (25). The repair of DNA damage in bacteria mainly relies on the homologous recombination (HR) pathway, which is a more stable and precise method compared with NHEJ. Therefore, the repair template provided to the bacteria can trigger DNA homology directed repair.

Two 500 bp DNA fragments corresponding to upstream and downstream regions of the *upp* gene in the FJSWX38M7 genome were cloned and ligated by overlap extension PCR to generate a repair template. Since pNZ624-sgRNA2 had the highest genome cleavage efficiency in the self-targeting experiment, the repair template was ligated to pNZ624-sgRNA2 to generate pNZ624-Δ*upp*-sgRNA2, allowing the strain to repair DNA damage caused by Cas3 cleavage (Fig. 2A). Plasmid pNZ624-Δ*upp*-sgRNA2 was delivered into FJSWX38M7 by electrotransformation, followed by random picking of colonies that had grown on mMRS agar supplemented with chloramphenicol. The deletion event of the *upp* gene in FJSWX38M7 was verified using PCR analysis. One of the samples yielded a smaller PCR product (2351 bp). Sanger sequencing also showed that this sample had a 642 bp deletion as expected, compared to wild-type. Uracil phosphoribosyl-transferase (UPRT) can convert 5-fluorouracil (5-FU) into 5-fluorodeoxyuridine monophosphate (5-FdUMP), thereby inhibiting the activity of thymidine nucleotide synthase, leading to cell death (26). Therefore, strains harboring an intact and expressed *upp* gene cannot grow in media supplemented with 5-FU. Overnight cultures of FJSWX38M7 wild-type and FJSWX38M7Δ*upp* were dropped onto mMRS agar supplemented with 5-FU. As expected, the strain with the deletion of the *upp* gene grew normally on agar containing 5-FU, whereas the wild-type failed to grow (Fig. 2C).

**Fig. 2.**
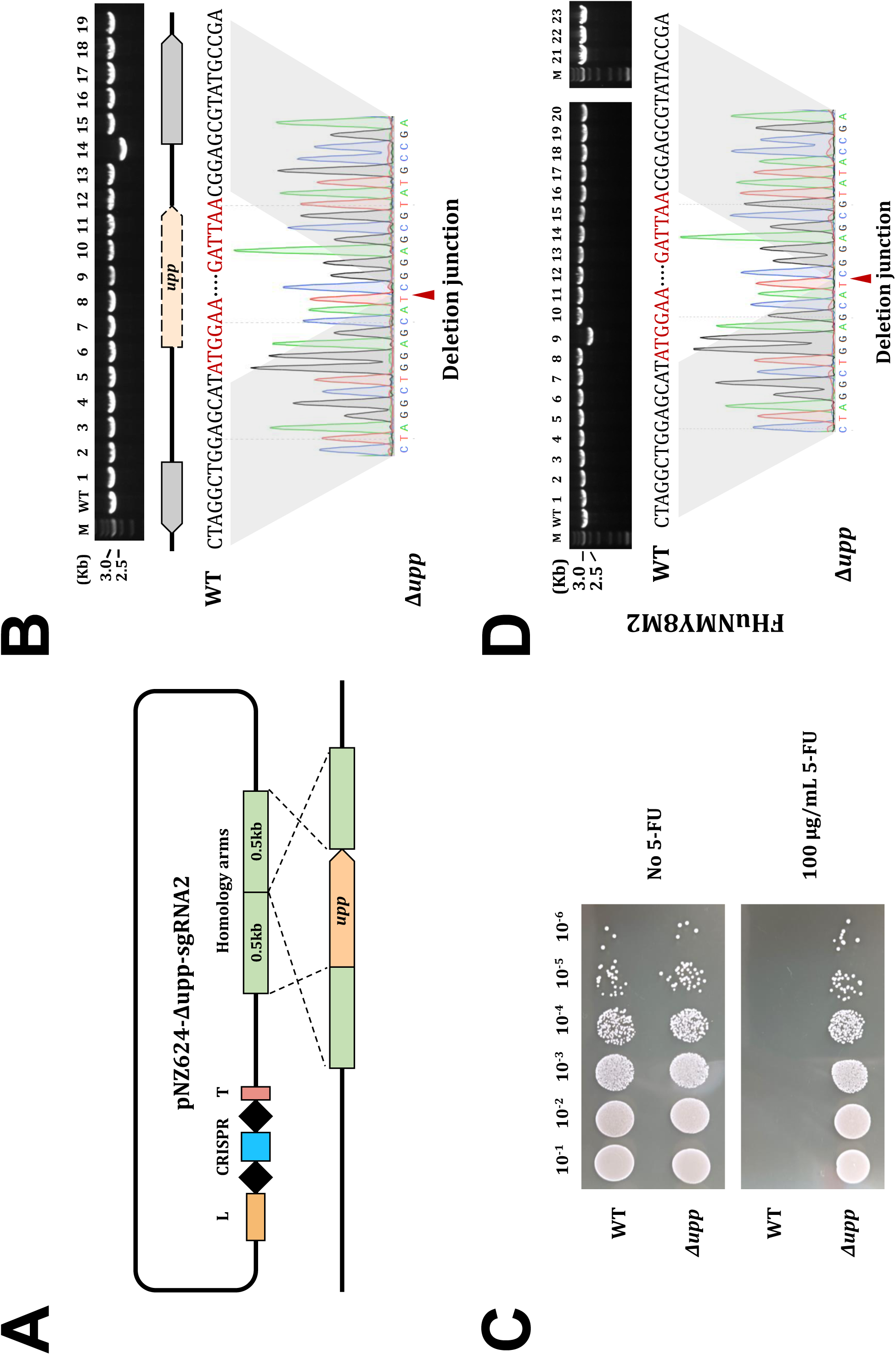
The *upp* gene deletion in *B. breve* using the endogenous Type I-C CRISPR-Cas system. (A) Schematic diagram of the *upp* gene deletion mediated by the endogenous I-C CRISPR-Cas system. The gene editing was constructed based on pNZ123, including a native leader, an artificial CRISPR array, a terminator, and a 1kb repair template. (B) Successful deletion of the *upp* gene in *B. breve* FJSWX38M7 was confirmed in one of the 19 randomly selected colonies by PCR and Sanger sequencing. M: marker; WT: wild-type. (C) The FJSWX38M7Δ*upp* mutant exhibited resistance to 100 μg/mL 5-FU while growing similarly on plates without 5-FU compared to the WT. Overnight liquid cultures of JSWX38M7 WT and Δ*upp* mutants were adjusted to the same OD_600_, then diluted to different concentrations, and 5μL of each dilution was spotted on plates with or without 5-FU. The plates were incubated at 37 °C for more than 36 h and then photographed. (D) PCR screening of Δ*upp* mutants in FHuNMY8M2. The *upp* deletion was observed in one out of 23 tested colonies. The chromatogram also confirmed the successful deletion of the *upp* gene by Sanger sequencing.

To explore whether genetic manipulation can also be performed in other *B. breve* strains that harbor the endogenous CRISPR-Cas system using this toolkit, *B. breve* FHuNMY8M2 was used as a target strain to accomplish the *upp* gene knockout. 23 Cm-resistant transformants were randomly selected, and PCR screening and sequencing confirmed that the *upp* gene was also successfully deleted in these derivatives of strain FHuNMY8M (Fig. 2D). These results demonstrated that the endogenous I-C CRISPR-based toolkit can be effectively used for genome engineering in *B. breve*.

### Precise point mutations and short DNA insertion in *B. breve* using the endogenous CRISPR gene editing toolkit

Point mutations occur widely during bacterial replication, including substitutions, insertions, and deletions of single bases. Point mutations can have a variety of deleterious or beneficial functional consequences, including changes in gene expression or alterations in encoded proteins (27). Point mutation of genes by genetic engineering technology is an important method to regulate gene expression, explore the key active sites of proteins, or change the activity of enzymes. To explore the potential of the CRISPR-based toolkit for precise point mutations, an sgRNA targeting the *upp* gene in strain FJSWX38M7 and a repair template containing a single-base substitution (193A>T) were cloned into plasmid pNZ624 (Fig. 3A). Forty transformants were screened by PCR, and Sanger sequencing results showed that the expected single base substitution appeared in 16 transformants, showing a high editing efficiency (Fig. 3C&D).

**Fig. 3.**
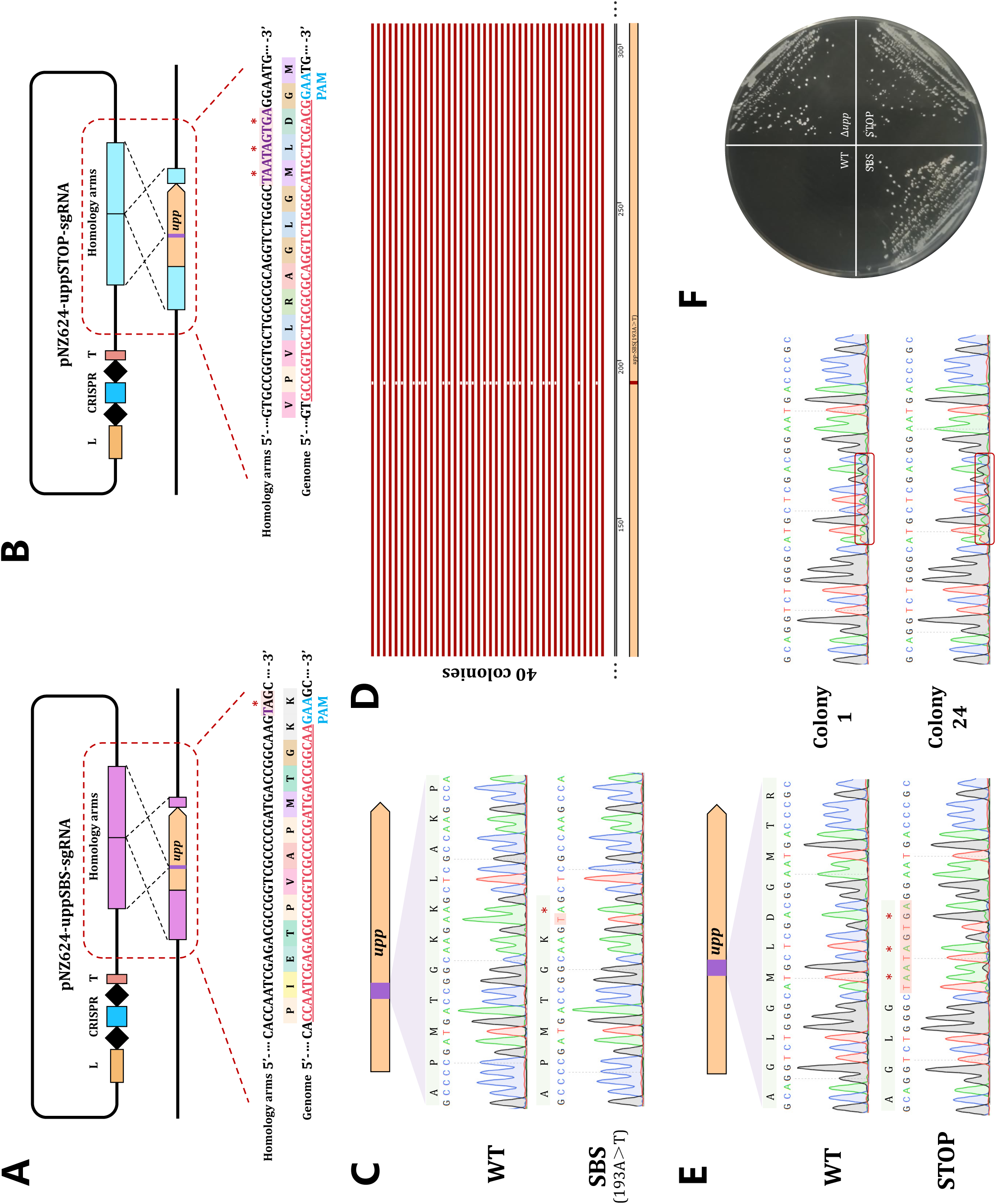
Single-base mutations and insertion of stop codons in *B. breve* using the endogenous CRISPR gene editing toolkit. (A) Schematic diagram of the single-base mutation mediated by the endogenous I-C CRISPR-Cas system. Single-base mutation sites were designed in the PAM region to prevent plasmid self-targeting. (B) Schematic diagram of stop codons insertion mediated by the endogenous I-C CRISPR-Cas system. Three consecutive stop codons were introduced in the *upp* gene by the repair template. (C) Sanger sequencing revealed that colonies on antibiotic plates carried 193A>T mutations in the *upp* gene that turned a lysine into a stop codon. (D) Multiple sequence alignment of Sanger sequencing results of 40 colonies was performed using Snapgene. The complete agreement with the reference sequence indicated that the colony with a single base mutation, while a base mismatch (white square) indicated that it was still wild-type. (E) The chromatogram revealed the successful insertion of three consecutive stop codons in the *upp* gene. The chromatograms of colonies 1 and 24 showed the stop codons insertion region with double peaks, indicating that they were mixed deletion mutants. (F) The FJSWX38M7 mutant with the single base mutation or insertion of stop codon could grow on the plate containing 100 μg/mL 5-FU, which proved that the translation of uracil phosphoribosyl-transferase was successfully terminated.

Gene insertion may allow strains to acquire new functions or completely disrupt gene function. In fact, horizontal gene transfer occurs frequently in bacteria and is an important driving force for their adaptation to the environment and evolution (28). To further expand *B. breve* genome editing capability, the short DNA insertion assay was performed in *B. breve* FJSWX38M7 using the endogenous I-C CRISPR-based toolkit. Triple stop codons were designed in the repair template and ligated into pNZ624-sgRNA2 to generate plasmid pNZ624-*upp*STOP-sgRNA, which is aimed at targeting the *upp* gene in FJSWX38M7 (Fig. 3B). Sanger sequencing demonstrated that the efficiency of stop codon insertion was 3/40, including a pure mutant and two mixed mutants (Fig. 3E).

In the single-base mutation assay, the lysine (**<u>A</u>**AG) of UPRT in FJSWX38M7 was mutated to a stop codon (**<u>T</u>**AG), resulting in the translation of UPRT being terminated (Fig. 3C). Similarly, the insertion of three consecutive stop codons into the *upp* gene in the short DNA insertion assay also caused premature termination of translation (Fig. 3E). The mutant strains FJSWX38M7-*upp*SBS and FJSWX38M7-*upp*STOP obtained from the above experiments were inoculated on plates supplemented with 5-FU and were shown to grow in the presence of this toxic uracil analogue as expected (Fig. 3F).

### Double gene deletion mediated by endogenous Type I-C CRISPR-Cas system in *B. breve*

In the study of gene clusters, it is often necessary to knock out different gene combinations to verify gene function, which requires gene editing tools to have the ability to edit multiple genes. For successive gene editing, the self-targeting plasmid must be cured from the edited cells before the next gene editing. To characterize the curing of the pNZ624-based gene editing plasmid, FJSWX38M7Δ*upp* harboring the plasmid was continuously cultured for nine passages in mMRS broth without adding antibiotics. Cultures were counted using the pour plate method in agar with or without antibiotics. Since cells without the plasmid cannot grow on the agar supplemented with chloramphenicol, the difference between counts on agar plates with and without antibiotics was assumed to represent the number of cells in which the plasmid had been cured. According to the enumeration results, the number of viable cells of FJSWX38M7Δ*upp* could reach ∼10^9^ CFU/mL stably in mMRS broth, but the number of cells containing the plasmid decreased by an order of magnitude every two generations (Fig. 4A). After one generation of culture in broth without antibiotics, the proportion of cells containing plasmids to the total number of cells dropped to 6%, and after three generations it was even lower than 0.5% (Fig. 4B). Hence, pNZ624-derived editing plasmids were easily cured by culturing in non-selective mMRS medium, ensuring the feasibility of successive gene editing in *B. breve*.

**Fig. 4.**
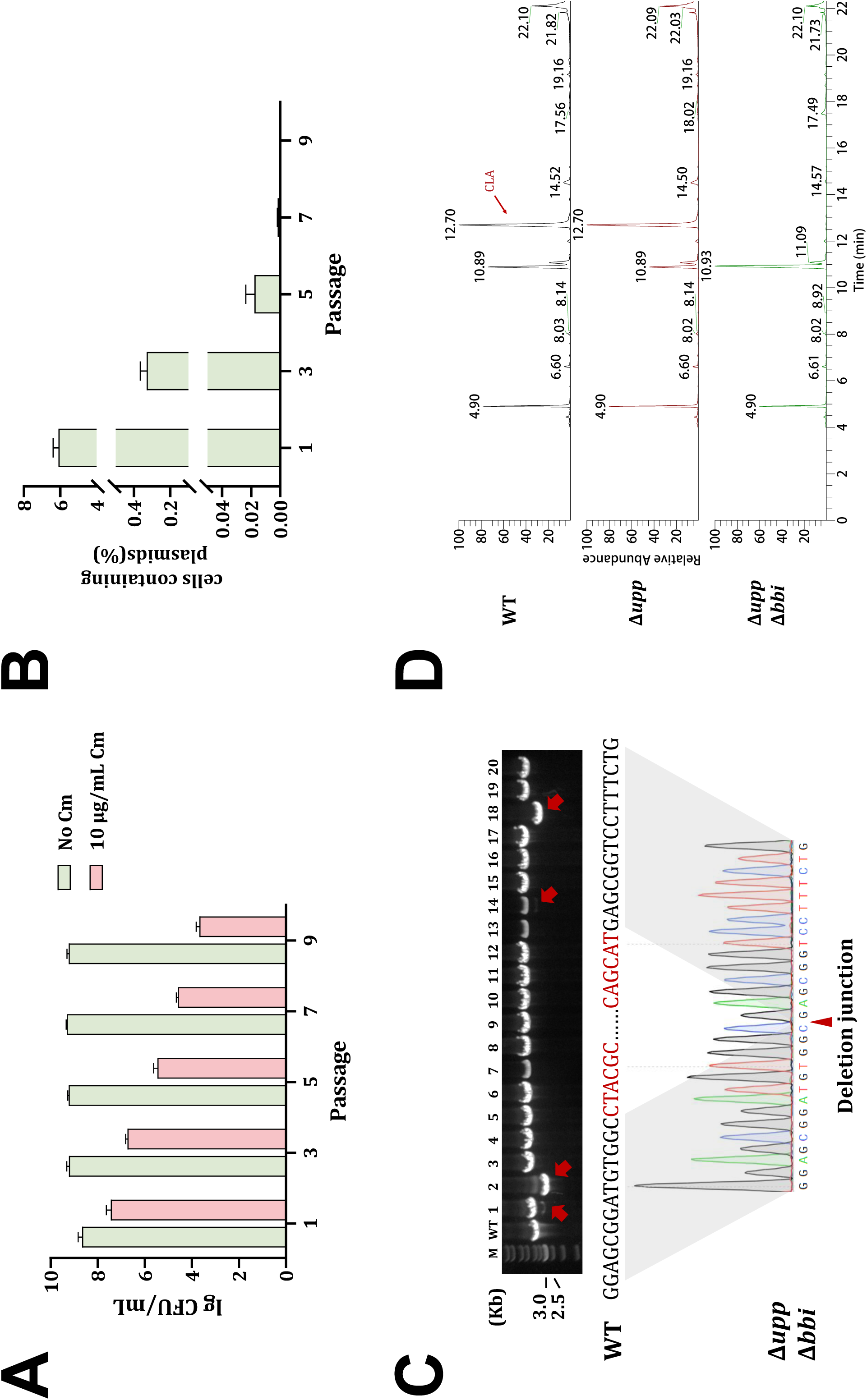
Successive gene editing in *B. breve* using the endogenous Type I-C CRISPR-Cas system. **(A)** Plasmid curing. FJSWX38M7Δ*upp* containing pNZ624-Δ*upp*-sgRNA2 was continuously cultured for nine generations in liquid medium without antibiotics, and the number of colonies that could grow on the antibiotic plate gradually decreased, which proved that the plasmid was gradually eliminated. (B) The proportion of cells containing plasmids to the total number of cells. (C) Successfully identified four *bbi* gene deletion mutants out of 20 randomly screened FJSWX38M7Δ*upp* transformants by PCR and Sanger sequencing, including 3 mixed mutants. (D) Gas chromatogram of conjugated linoleic acid production. The retention time of conjugated linoleic acid was 12.70 min.

To validate the gene knock system beyond the *upp* gene and to establish if a strain with multiple successive gene knock outs can be constructed, we next targeted the *bbi* gene. Conjugated linoleic acid (CLA) is a functional fatty acid widely used in food, and *B. breve* is the species with strong biotransformation of CLA among bifidobacteria (29). The *bbi* gene in *B. breve* encodes a linoleic acid isomerase that directly converts the substrate linoleic acid (LA) to CLA in a one-step reaction without other intermediates (30). Therefore, the *bbi* gene in FJSWX38M7Δ*upp* was selected as the next target gene. pNZ624-Δ*bbi*-sgRNA was introduced into the plasmid-cured FJSWX38M7Δ*upp* to generate a strain with the double deletion of the *upp* gene and the *bbi* gene. PCR analysis showed that colonies 1, 2, 14, and 18 carried a 993 bp deletion, showing an editing efficiency of 20% (4/20) (Fig. 4C). The CLA transformation ability of mutants was analyzed using GC-MS. The results showed that the wild-type or FJSWX38M7Δ*upp* had the ability to convert LA into CLA, while the presence of CLA was not detected in the fermentation broth of FJSWX38M7Δ*upp*Δ*bbi* (Fig. 4D). These results demonstrate that successive gene editing of *B. breve* is feasible based on its endogenous I-C CRISPR-Cas system.

## Discussion

*B. breve* is one of the dominant species in the gut of breast-fed newborns and is important for maintaining health in early life (3). Evidence from animal model studies and human clinical trials had previously shown that *B. breve* plays an important role in the prevention and treatment of gastrointestinal diseases (31), neurological diseases (32) and allergic diseases (33). However, the lack of genetic tools and highly active restriction modification (R-M) systems hampered the investigation of the molecular mechanisms behind their purported benefits (4, 34). It has been reported that pORI19 lacking a *repA* gene can be used to accomplish insertional mutagenesis in *B. breve* (35). Overcoming the barriers of the R-M system by artificially removing restriction sites in the plasmid or by artificially methylating the plasmid has proven effective (36, 37). However, the non-replicative plasmids used in these strategies will introduce antibiotic resistance genes into the *B. breve* genome, which brings difficulties to successive gene editing and the risks of horizontal transfer of resistance genes. In comparison, editing methods based on a CRISPR-Cas system can achieve efficient and seamless deletion of genes. A recent study showed that the exogenous CRISPR-Cas9-based editing system can be successfully applied to gene knockout in *B. animalis* AR668 (38). Another study restructured the endogenous I-G type system and introduced the exogenous base editor in *B. animalis* subsp*. lactis* to achieve gene deletion and single-base mutation, providing a solid basis for attempting a similar editing strategy based on the CRISPR-Cas system in *B. breve* (20).

The widespread presence of an endogenous CRISPR-Cas system in the *B. breve* genome presented this system as an attractive opportunity for gene editing. Type I-C is the predominant subtype in *B. breve*, and relies on the degradation of the non-target strand by the nuclease Cas3 to form a DNA nick (39). In fact, DNA double-strand breaks caused by Cas9 may be irreparable by the host (40), and even the expression of Cas9 was strongly toxic in some strains (16, 17). The advantage of the endogenous CRISPR-Cas system editing strategy is that only the sgRNA and the repair template need to be provided by the plasmid since the nuclease is expressed by the host. This can greatly reduce the length of the plasmid, which is conducive to its efficient delivery and associated circumvention of R-M systems.

Plasmid pNZ123 was used as the basis for gene editing vectors and was shown to be efficiently electro-transformed into *B. breve* (36). Plasmid interference assay demonstrated the activity of the endogenous I-C CRISPR-Cas system in FJSWX38M7, and then self-targeting experiments were designed to explore the feasibility of gene editing based on the endogenous I-C CRISPR-Cas system. A native leader and an artificial CRISPR array were used to transcribe crRNA to better mimic the expression and maturation stages of the CRISPR-Cas system. The non-essential gene *upp* encoding UPRT, was selected as the target gene since the deletion mutants of *upp* can easily be distinguished from the wild-type by growing on agar supplemented with 5-FU (41). Three different sgRNAs showed different cleavage efficiencies, similar phenomena are commonly observed in gene editing of animals and plants based on CRISPR-Cas9, which may be related to the GC content of the sgRNA and the location of the targeted gene (42, 43). The *upp* gene deletion was successfully achieved in two different *B. breve* strains using pNZ624-sgRNA2 supplemented with the repair templates. But escaping from CRISPR targeting was also observed. Escape targeting may not be related to the mutations of PAM-protospacer regions in the genome (18). The reasons for the escape transformants were speculated to be as follows: 1) Surviving transformants escaped CRISPR targeting due to the mutation or deletion of *cas* operators (44, 45). 2) Low gene editing efficiency was due to the genetic recalcitrance of *B. breve* (46). 3) This may be attributed to insufficient expression of cascade and/or sgRNA (40).

Due to the lack of NHEJ repair mechanisms, it is necessary to introduce repair templates in *B. breve*. The mutated site and inserted sequence can be designed in the repair template to achieve gene mutation and seamless insertion. Here, point mutations and short DNA insertions in the *upp* gene were successfully achieved using the endogenous CRISPR gene editing toolkit, and the gene point mutation showed high efficiency. Cytosine base editors (CBEs) and adenine base editors (ABEs) are commonly used for base editing in animal and plant cells. Taking CBE as an example, it converts the cytosine of the target DNA into uracil through the fusion expression of cytidine deaminase and inactive dCas9, which can convert the cytosine of the target DNA into uracil, which is further repaired by cells into thymine, resulting in a change from C to T. To improve base editing efficiency, Cas9 nickase (D10A) can be used instead of dCas9 to force cells to use deaminated DNA strands as templates (47). But its disadvantage is that the mutation from C to T will be formed immediately at the DNA target region, and the desired mutation needs to be screened later. Base editing using CBE has been achieved in some bacterial species (20, 48).

The pNZ624-based gene editing plasmids were easily cured by continuous culture in medium (both broth and plate) without supplementation of antibiotics, which ensured their feasibility for successive genome editing in *B. breve*. A commonly used plasmid curing method is to connect the sucrose sensitive gene *sacB* to the plasmid, and the survivors on the sucrose plate are the colonies without the plasmid (48). However, this method is not suitable for *B. breve*, because the preparation of electroporation competent cells of *B. breve* needs to be preserved in sucrose ammonium citrate solution. Another approach is to use plasmids with thermosensitive replicons, but given the lack of genetic tools in bifidobacteria, more plasmids that can be used in bifidobacteria need to be developed. A recent study provided a strategy for plasmid elimination using a lactose-induced second sgRNA targeting the resistance gene of this plasmid (38). Cas9 was guided by the first sgRNA to cleave the target gene when lactose was not added. The addition of lactose-induced transcription of the second sgRNA directed Cas9 to cleave the plasmid itself when target gene deletion was complete.

In this study, the gene editing efficiency based on the endogenous I-C CRISPR-Cas system in *B. breve* was similar to that implemented using endogenous I-G CRISPR-Cas system in *B. animalis* subsp*. lactis*, and there is still further room for improvement, such as: 1) Construction of dual sgRNAs. Since Cas3 will cut non-target DNA strands, using sgRNAs targeting the coding strand and the non-coding strand separately will cause DNA double-strand breaks and reduce escape targeting. 2) Changing the length of the repair template. Studies have shown that an appropriate repair template length was beneficial to improve editing efficiency (18, 38). 3) Introducing the auxiliary repair system. Recombinase systems, such as lambda Red, RecAb, and Rac-RecET, can improve the homologous recombination efficiency of repairing templates and genomes to increase the positive rate of transformants (48).

Overall, this study successfully achieved gene deletion, point mutation, gene insertion, and successive gene editing in *B. breve* using the endogenous CRISPR gene editing toolkit. It represents the first report of gene editing in *B. breve* based on an autochthonous CRISPR-Cas system, providing a new tool for functional gene mining in *B. breve* and opening up opportunities for the development of engineered biotherapeutics through genetic engineering.

## Authors and contributors

**Xiao Han**: Methodology, Validation, Investigation, Data curation, Writing-original draft, Visualisation. **Lulu Chang**: Writing-review & editing. **Haiqin Chen**: Writing-review & editing. **Jianxin Zhao**: Resources, Data curation. **Fengwei Tian**: Writing-review & editing, Conceptualisation, Supervision, Funding. **R. Paul Ross**: Writing-review & editing. **Catherine Stanton**: Writing-review & editing. **Douwe van Sinderen**: Writing-review & editing. **Wei Chen**: Conceptualization, Supervision, Project administration, Funding. **Bo Yang**: Conceptualization, Validation, Investigation, Data curation, Writing-original draft, Writing-review & editing, Visualization, Supervision.

## Methods

### Bacterial strains, plasmids, oligonucleotides and culture conditions

The bacterial strains and plasmids used in this study are listed in Table S1, and the oligonucleotides used are listed in Table S2. *E. coli* was cultured on Luria-Bertani (LB) agar or in LB broth at 37°C with shaking. *B. breve* was propagated anaerobically (80% N_2_, 10% CO_2_, 10% H_2_) at 37°C on modified de Man Rogosa Sharpe (mMRS) agar or in mMRS broth. The composition of mMRS was as follows: 10 g/L tryptone, 10 g/L beef extract, 5 g/L yeast extract, 20 g/L D-(+)-glucose, 2.6 g/L K_2_HPO_4_, 2 g/L diammonium hydrogen citrate, 2 g/L sodium acetate, 0.1 g/L MgSO_4_·7H_2_O, 0.05 g/L MnSO_4_·H_2_O, 1 g/L Tween80, 1 g/L L-cysteine. If necessary, 10 μg/mL chloramphenicol was supplemented to the medium for selecting *E. coli* recombinants or *B. breve* mutants. 5-FU (final concentration 100 μg/mL) was added to mMRS agar to culture *B. breve* Δ*upp* mutants.

### DNA manipulations

*B. breve* genomic DNA was extracted using the TIANamp Bacteria DNA Kit (TIANGEN, Beijing, China), and plasmid DNA was extracted with the TIANprep Mini Plasmid Kit (TIANGEN, Beijing, China) according to the instructions of the manufacturer. The oligonucleotides used in this study were synthesized by Shanghai Sangon. DNA fragments were amplified using 2 × Phanta Flash Master Mix (Vazyme, Nanjing, China) following standard PCR procedures. PCR products were analyzed on 1% agarose gels, and DNA fragments were purified using the GeneJET Gel Extraction Kit (Thermo Scientific, Massachusetts, USA). The digested plasmid and DNA fragment (if necessary) were ligated with T4 DNA ligase (Thermo Scientific) or One Step Cloning Kit (YEASEN, Shanghai, China) and then transformed into *E. coli*. The correct recombinants were confirmed by PCR using specific primers. Sanger sequencing service was provided by Azenta (Suzhou, China).

Oligonucleotides were phosphorylated and annealed as previously described (49). Briefly, paired oligonucleotides were incubated at 37°C for 30 min in a reaction mixture supplemented with T4 PNK (NEB, Massachusetts, USA) and T4 Ligation Buffer (NEB, Massachusetts, USA). The reaction mixture was incubated at 95°C for 5 min and then ramped down to room temperature at 5°C/min. Diluted annealed oligonucleotides and digested plasmids were ligated overnight at 16°C using T4 DNA ligase and then transformed into *E. coli*.

### Construction of plasmids

Plasmid pNZ123 is a replication shuttle vector for *E. coli* and bifidobacteria. Diluted annealed S82-123-F/S82-123-R or PS82-123-F/PS82-123-R and XhoI-EcoRI digested pNZ123 were ligated to obtain interference plasmids pNZ123-S82 or pNZ123-PS82. Endogenous CRISPR-based editing vector pNZ624 was obtained by ligation of artificial crRNA synthesized from GenScript and NspI-digested pNZ123, wherein the artificial crRNA included a leader sequence, a sgRNA insertion site, two repeats, and a rho-terminator. The annealed oligonucleotides uppsgRNA1-F/R were ligated with BsaI-digested pNZ624 to obtain pNZ624-sgRNA1 for self-targeting experiments. pNZ624-sgRNA2 and pNZ624-sgRNA3 were also obtained using the method described above. The upstream and downstream homology arms of the *upp* gene in FJSWX38M7 were amplified using primers 21uppHAL-F/R and 21uppHAR-F/R, respectively. The two purified fragments were ligated by overlap extension PCR to obtain a 1 kb repair template. Both the repair template and pNZ624-sgRNA2 were digested by XhoI-EcoRI and ligated to obtain the *upp* gene knockout plasmid pNZ624-Δ*upp*-sgRNA2. The FHuNMY8M2 *upp* gene knockout plasmid pNZ624-Δ*upp*-sgRNA16 was also constructed with pNZ624-sgRNA2 as the backbone, using primers 16uppHAL-F/R and 16uppHAR-F/R by the same method.

In order to obtain plasmid pNZ624-*upp*SBS-sgRNA, primers 21HALSBS-F/R and 21HARSBS-F/R were used to amplify the upstream and downstream homology arms, and the mutated bases were introduced by primers 21HALSBS-R and 21HARSBS-F. The upstream and downstream homology arms were ligated by overlap extension PCR to generate a repair template containing single-base substitutions. Finally, the repair template and the XhoI-EcoRI-digested plasmid containing sgRNA (BsaI-digested pNZ624 ligated with annealed uppsgRNASBS-F/R) were ligated to obtain the single-base mutant plasmid. Similarly, homology arms designed to introduce three stop codons in the *upp* gene were amplified by PCR using the primers 21HALSTOP-F/R and 21HARSTOP-F/R. Overlap extension PCR amplified repair template was cloned into XhoI-EcoRI-digested pNZ624-sgRNA2 to generate plasmid pNZ624-*upp*STOP-sgRNA. The construction method of *bbi* gene knockout plasmid pNZ624-Δ*bbi*-sgRNA was the same as that of pNZ624-Δ*upp*-sgRNA2, and the oligonucleotides used in this process were bbisgRNA-F/R, 21bbiHAL-F/R, and 21bbiHAR-F/R.

### Preparation of *E. coli* competent cells and transformation

A fresh 37°C overnight culture of *E. coli* TOP10 was inoculated into 50 mL of LB broth. The culture was grown at 37°C with shaking (200 rpm) until OD_600_ reached 0.5. And then the culture was frozen on ice for 10 min. The cells were harvested at 4000 rpm for 10 minutes at 4°C. The cell pellet was resuspended in 10 mL of 0.1mol/L CaCl_2_ solution and frozen on ice for 20 min, followed by cell harvetsing by centrifugation (4000 rpm, 10 minutes at 4°C). The cell pellet was again resuspended in CaCl_2_ and chilled on ice, followed by centrifugation steps. Finally, the pellet was resuspended in 1 mL of 0.1 mol/L CaCl_2_ glycerol solution (1 mol/L CaCl_2_ solution and 22% glycerin solution were mixed at a volume ratio of 1:9) and aliquoted in 100 µL volumes into pre-cooled 1.5 mL centrifuge tubes and stored at -80°C until use.

10 μL of the ligation product or 200 ng of the plasmid were added to 100 μL of *E. coli* competent cells, mixed by flicking, and incubated on ice for 30 minutes. The mixture was heated at 42 °C for 90 s and frozen again on ice for 2 min. 890 μL (preheated at 37°C) of LB broth was added to the mixture and incubated at 37°C with shaking (200 rpm) for 60 minutes. An appropriate volume of cultures was spread on selective LB agar and incubated overnight at 37°C. Primers pNZ123-F/R were used for PCR screening of recombinant plasmids to confirm that the gene fragments and vectors were correctly ligated.

### Preparation of *Bifidobacterium* competent cells and transformation

Preparation and transformation of of competent *B. breve* cells was based on a previously described protocol with modifications (36). A fresh 5 mL overnight culture of *B. breve* was inoculated into 45 mL of mMRS broth and grown anaerobically at 37°C until the OD_600_ reached between 0.6 and 0.9. After the culture was chilled on ice for 20 min, the cells were harvested by centrifugation (6000 rpm, 10 minutes at 4°C). The cell pellet was washed twice with pre-cooled sucrose citrate buffer (0.5M sucrose and 1mM ammonium citrate, pH 5.8) under the same centrifugation conditions. The final harvested cell pellet was resuspended in 200 μL ice-cold sucrose-citrate buffer and distributed according to the volume of 50 μL per tube. The competent cells of *B. breve* were all freshly prepared because the transformation efficiency was shown to decrease significantly following cold storage.

200 ng of plasmid was added to 50 μL *B. breve* competent cells, flicked to mix well, and then transferred to a pre-cooled 2 mm electroporation cuvette. The electroporation cuvette was pulsed with 2000 V using an Eppendorf electroporator. After electrotransformation, 950 μL mMRS broth (preheated at 37°C) was added to the cells and incubated at 37°C for 2.5 hours under anaerobic conditions. An appropriate volume of culture was plated on selective mMRS agar and grown anaerobically at 37°C for 36-72 hours. Primers used to identify mutations or deletions in the obtained transformants were listed in Table S2.

### Plasmid curing

FJSWX38M7Δ*upp* containing the plasmid was inoculated at 2% (vol/vol) in the mMRS broth without antibiotics, cultured anaerobically at 37°C for 24 hours, and then inoculated the next generation by the same operation, repeated nine times. The cultures of each passage were counted using the pour plate method in medium with or without antibiotics. The cure ratio of the plasmid was recorded as the difference between the counts of the two plates divided by the counts of the non-selective plates.

*B. breve* recombinants containing the plasmid were cultured in mMRS broth without antibiotics for two passages and then streaked on mMRS agar without antibiotics. Single colonies were streaked onto mMRS agar with or without antibiotics. If the colony cannot grow on the antibiotic agar, it indicated that its plasmid has been cured.

### Extraction and analysis of conjugated linoleic acid (CLA)

CLA in the culture was extracted and analyzed as previously described (50). Briefly, *B. breve* was cultured for 72 h in mMRS broth containing 0.5 mg/mL linoleic acid. 2 mL of isopropanol was added to 3 mL of culture and mixed by vigorous shaking. 3 mL of hexane was added to the mixture and shaken vigorously. After standing, the supernatant was transferred to a clean glass bottle and dried using the nitrogen blower. Then 400 μL of methanol was added to the glass bottle and shaken thoroughly. An appropriate amount of trimethylsilylated diazomethane was added to the bottle until the yellow color of the solution remained unchanged for 15 min, indicating that the methyl esterification was complete. The above samples were dried again under nitrogen. 1mL of hexane was added to the glass bottle, shaken thoroughly and then centrifuged at 5000 rpm for 5 min. 800 μL of the supernatant was transferred to a 2 mL vial for detection by GCMS-QP2010 Ultra (Thermo Scientific, Massachusetts, USA). The specific parameters were as follows. A gas phase column RT-5MS (30 m × 0.25 mm × 0.25 μm) was used. The initial temperature of the program was 180 °C and kept for 3 min, then raised to 190 °C at a rate of 10 °C/min and kept for 3 min. Then the temperature was raised to 220°C at 5°C/min and kept for 1 min. Finally, the temperature was raised to 230°C at a rate of 2°C/min. The temperature of the injection port and the interface were both 240°C and the temperature of the ion source was 230°C.

### Statistical analysis

Statistical analysis was performed using an unpaired two-tailed Student’s t-test. \**P*<0.05 and \*\**P*<0.01 were considered statistically significant.

## Supplementary figures

**Fig. S1 Derived model for type I-C cascade-mediated DNA interference.** Rose red: Cas5; Orange: Cas8; Blue: Cas7; Green: Cas3.

**Fig. S2 Occurrence frequency of Type I-C PAM sequences in the *B. breve* FJSWX38M7 genome.** The horizontal axis is the distance between two PAMs, defined as the number of nucleotides from the previous 5’-TTC-3’ or 5’-GAA-3’ to the next 5’-TTC-3’ or 5’-GAA-3’. The vertical axis indicates the number of each distance.

**Fig. S3 Leader sequence of Type I-C CRISPR array in *B. breve* using the WebLogo server.** The relative frequency of each nucleic acid at that position in the 250 bp leader sequence is displayed. Strains with interrupted leader sequences by mobile elements were removed by manual screening.

**Fig. S4 Construction steps of the gene editing vector based on pNZ624.** pNZ624 was obtained by ligation of NspI digested pNZ123 and an artificial crRNA, including a leader sequence, a sgRNA insertion site, two repeats and a rho-terminator. The sgRNA insertion site contains two BasI restriction sites with opposite sequences for easy insertion of sgRNA. Phosphorylated and annealed synthetic oligos were ligated into BsaI-digested pNZ624 to generate vectors containing sgRNA. Finally, the left and right repair templates of the target gene were ligated into the vector containing sgRNA to generate the gene editing plasmid.

**Fig. S5 The number of different spacer lengths in *B. breve*.**

